# Diurnal rhythm causes metabolic crises in the cyanobacterial mutants of c-di-AMP signalling cascade

**DOI:** 10.1101/2023.11.14.567006

**Authors:** Michael Haffner, Oliver Mantovani, Philipp Spät, Boris Maček, Martin Hagemann, Karl Forchhammer, Khaled A. Selim

## Abstract

In nature, the photoautotrophic lifestyle of cyanobacteria has to cope with the successive diurnal changes in light supply. Light supply throughout the day enables photosynthesis and glycogen biosynthesis, while night phases require the switch to a heterotrophic-like lifestyle relying on glycogen catabolism. We previously highlighted a unique function of the carbon control protein, SbtB, and its effector molecule c-di-AMP, for the nighttime survival of cyanobacteria through the regulation of glycogen anabolism. However, the extent to which c-di-AMP and SbtB impact the cellular metabolism for day-night survivability remained elusive. To gain better understanding of cellular processes regulated by SbtB or c-di-AMP, we compared the metabolomic and proteomic landscapes of Δ*sbtB* and the c-di-AMP-free (Δ*dacA*) mutants of the model strain *Synechocystis* sp. PCC 6803. While our results indicate that the cellular role of SbtB is restricted to carbon/glycogen metabolism, the diurnal lethality of Δ*dacA* seems to be a sum of dysregulation of multiple metabolic processes. These processes include photosynthesis and redox regulation, which lead to elevated levels of intracellular ROS and glutathione. Further, we show an impact of c-di-AMP on central carbon as well as on nitrogen metabolism. Effects on nitrogen metabolism are linked to reduced levels of the global nitrogen transcription regulator NtcA and highlighted by an imbalance of the glutamine to glutamate ratio as well as reduced metabolite levels of the arginine pathway. We further identified the HCO_3_^-^ uptake systems, BicA and BCT1 as novel SbtB targets, in agreement with its broader role in regulating carbon homeostasis.

## Introduction

About 2.7 to 3.2 billion years ago, the evolution of two coupled photosystems in primordial cyanobacteria permitted water splitting and created the first aerobic life in Earth’s history (Knoll 2008). Fundamental to the autotrophic life-style is the tight coupling between oxygenic photosynthesis and inorganic carbon (Ci; referring to CO_2_ or HCO_3_^−^) fixation via the Calvin-Benson-Bassham cycle, using energy provided by sunlight throughout the day. On Earth, however, the ambient light supply oscillates through successive day-night cycles. Therefore, night phases require a metabolic switch to heterotrophic-like carbon catabolism, which includes glycogen catabolism mainly via the oxidative pentose-phosphate (OPP) pathway (Gründel et al. 2012, Diamond et al. 2015, Welkie et al. 2018, Makowka et al. 2020). Confronted with this perpetuating environmental cue, cyanobacteria evolved regulatory processes to adapt their metabolism to the natural diurnal rhythm. These involve the sensing of intracellular carbon and nitrogen levels as well as the sensing of the cellular redox and energy status (Diamond et al. 2017, Köbler et al. 2018, Welkie et al. 2018, Lucius et al. 2022). Furthermore, cyanobacteria evolved an intricate circadian clock to sense and anticipate to consecutive day-night cycles (Welkie et al. 2019). In addition, the second messenger nucleotides guanosine penta- and tetraphosphate [(p)ppGpp] as well as cyclic di-adenosine monophosphate (3′,5′-c-di–adenosine 5′-monophosphate; hereafter c-di-AMP) have recently been shown to be involved in regulating cyanobacterial diurnal lifestyle (Hood et al. 2016, Puszynska and O’Shea 2017, Rubin et al. 2018, Mantovani et al. 2023a). The latter by regulation glycogen synthesis via the PII-like protein SbtB (Selim et al. 2021).

SbtB, a small regulatory protein belonging to the PII signaling superfamily (Forchhammer et al. 2022), is encoded in an operon together with the low Ci-induced sodium-dependent bicarbonate transporter SbtA (Selim et al. 2018, Forchhammer and Selim 2020). Similar to canonical PII proteins (Lapina et al. 2018, Selim et al. 2019 & 2020a), SbtB binds the adenine nucleotides ATP and ADP, but in contrast to these, it also binds AMP, cAMP and c-di-AMP (Selim et al. 2018 & 2021), thus combining the cell’s energy status with second messenger signaling, which directly influences the SbtA-B complex formation (Selim et al. 2023, Haffner et al. 2023) and many other processes related to Ci acclimation (Mantovani et al. 2022, 2023a & 2023b). Under low Ci conditions, intracellular AMP levels rise, thus leading to a strong SbtA-B complex in which SbtB serves as plug-like structure to prevent HCO_3_^-^ leakage (Haffner et al. 2023, Selim et al. 2023). Under high Ci conditions, which are sensed by high intracellular levels of ATP and cAMP, SbtB partially dissociates from SbtA (Selim et al. 2018 & 2023).

The transcriptomic analysis of a SbtB knockout mutant (Δ*sbtB*) revealed a broader regulatory impact of SbtB on cyanobacterial Ci acclimation that is not solely restricted to the regulation of SbtA (Mantovani et al. 2022). Deletion of SbtB induces a low Ci (LC) pre-acclimated state, while only a few genes that are directly involved in the carbon-concentrating mechanism (CCM) are differently expressed in a Δ*sbtB* mutant as compared to *Synechocystis* wild type (WT). Therefore, an interaction of the PII-like SbtB protein with yet unknown factors, similar to what is known for the versatile canonical PII proteins is hypothesized (Forchhammer et al. 2022, Mantovani et al. 2022). Notably, SbtB has an effect on the expression of several transcription factors (Mantovani et al. 2022), among them a major regulator of Ci acclimation NdhR. Moreover, the imbalance in Ci acclimation in Δ*sbtB* mutant has further effects on the expression of genes involved in nitrogen metabolism, which is reflected by an imbalanced glutamate to glutamine ratio.

Recently, c-di-AMP, one of the newly discovered second messengers, turned out to play important roles in cyanobacterial acclimation mechanisms (Mantovani et al. 2023a). In Firmicutes, c-di-AMP is known to control cell wall integrity and osmotic homeostasis by regulating osmolyte and K^+^ and Mg^2+^ ion transporters (Stülke and Krüger 2020, Yin et al. 2020). Identification of c-di-AMP target proteins in the cyanobacterium *Synechocystis* sp. PCC 6803 (hereafter *Synechocystis*) revealed several ion transporters like in firmicutes (Selim et al. 2021). These findings support an involvement of c-di-AMP in cyanobacterial osmoregulation and fit to the fluctuating c-di-AMP levels among abiotic stress (Agostoni et al. 2018). Concomitantly, the c-di-AMP deficient *Synechocystis* mutant (Δ*dacA*), is lethal under osmotic stress, generated by high amounts of extracellular sugars (Selim et al. 2021). Unique to cyanobacteria, c-di-AMP regulates additionally the nocturnal dormancy, as the c-di-AMP deficient mutants of *Synechococcus elongatus* were impaired in diurnal growth, which was recently confirmed in *Synechocystis* as well (Rubin et al. 2018, Selim et al. 2021).

The PII-like SbtB protein was identified as major receptor of the c-di-AMP messenger in cyanobacteria. In its c-di-AMP bound state, SbtB was shown to regulate glycogen synthesis by interacting with the glycogen branching enzyme (GlgB) during the day-phases (Selim et al. 2021), and therefore both Δ*sbtB* and Δ*dacA* are not able to accumulate sufficient glycogen amount during the day phase. As glycogen catabolism via the OPP pathway (Welkie et al. 2018) is required for nighttime survival, the regulation of diurnal glycogen synthesis by the SbtB-c-di-AMP complex represents an important level of controlling day-night cycles. The broad impact of *sbtB* on cyanobacterial physiology, as well as the wide range of putative c-di-AMP receptors indicate that these factors are not solely restricted to the regulation of glycogen synthesis. In this study, we used a targeted-metabolomics approach and correlated it to the proteomics landscape of Δ*sbtB* and Δ*dacA* mutants in comparison to *Synechocystis* WT throughout diurnal growth, to get further insights into the regulatory modes of both SbtB and c-di-AMP on diurnal metabolism.

## Results & Discussion

### Targeted metabolomics reveals metabolic dysregulation in direct response to the shift from constant light to diurnal light/ dark cycles

Our previous physiological experiments revealed that both *Synechocystis* mutants, the c-di-AMP biosynthesis mutant (Δ*dacA*) and the c-di-AMP receptor mutant (Δ*sbtB*) are deficient in diurnal glycogen accumulation and show poor growth under diurnal (day-night) conditions (Selim et al. 2021). This deficiency was attributed to the less than 30% glycogen content in the Δ*dacA* and Δ*sbtB* mutants relative to WT and concomitant energy crisis in the night period. Accordingly, we expected characteristic metabolic changes in central carbon and the closely related nitrogen metabolism. Giving their common function in glycogen accumulation, we assumed a partial overlap of their metabolic profiles during the night when carbon resources become limiting upon transition from photoautotrophic to heterotrophic metabolism. Furthermore, we expected the metabolic defects to aggravate following the initial shift from growth-permissive, constant illumination to subsequent alternating 12 h day/12 h night cultivation. In the course of these cycles, the mutations should increasingly affect metabolism until the capability of night-time survival is lost as described previously (Selim et al. 2021).

To test these expectations and to characterize the impact of the Δ*dacA* and Δ*sbtB* mutations on the primary metabolism, we used targeted LC-MS based metabolomics analyses that compared quantitatively key central metabolites in WT and mutants by time resolved analysis of 12 h day/12 h night cycles after two days acclimation of the cultures to the diurnal rhythm. Samples were taken at day 3 after 11 h illumination, 1 h after onset of darkness, 4 h darkness, 11 h darkness, and at the following day at 1 h and 11 h after illumination (**Fig. 1, Supplemental Fig. S1 and Table S1**). Statistical analysis revealed that the metabolome of Δ*dacA* cells deviated strongly from WT, whereas Δ*sbtB* cells had an intermediate metabolic phenotype (**Supplemental Fig. S2 and Table S2**) supporting our assumption of a partial overlap, and furthermore, points at a broader impact of c-di-AMP signaling, independent from SbtB.

**Fig. 1:**
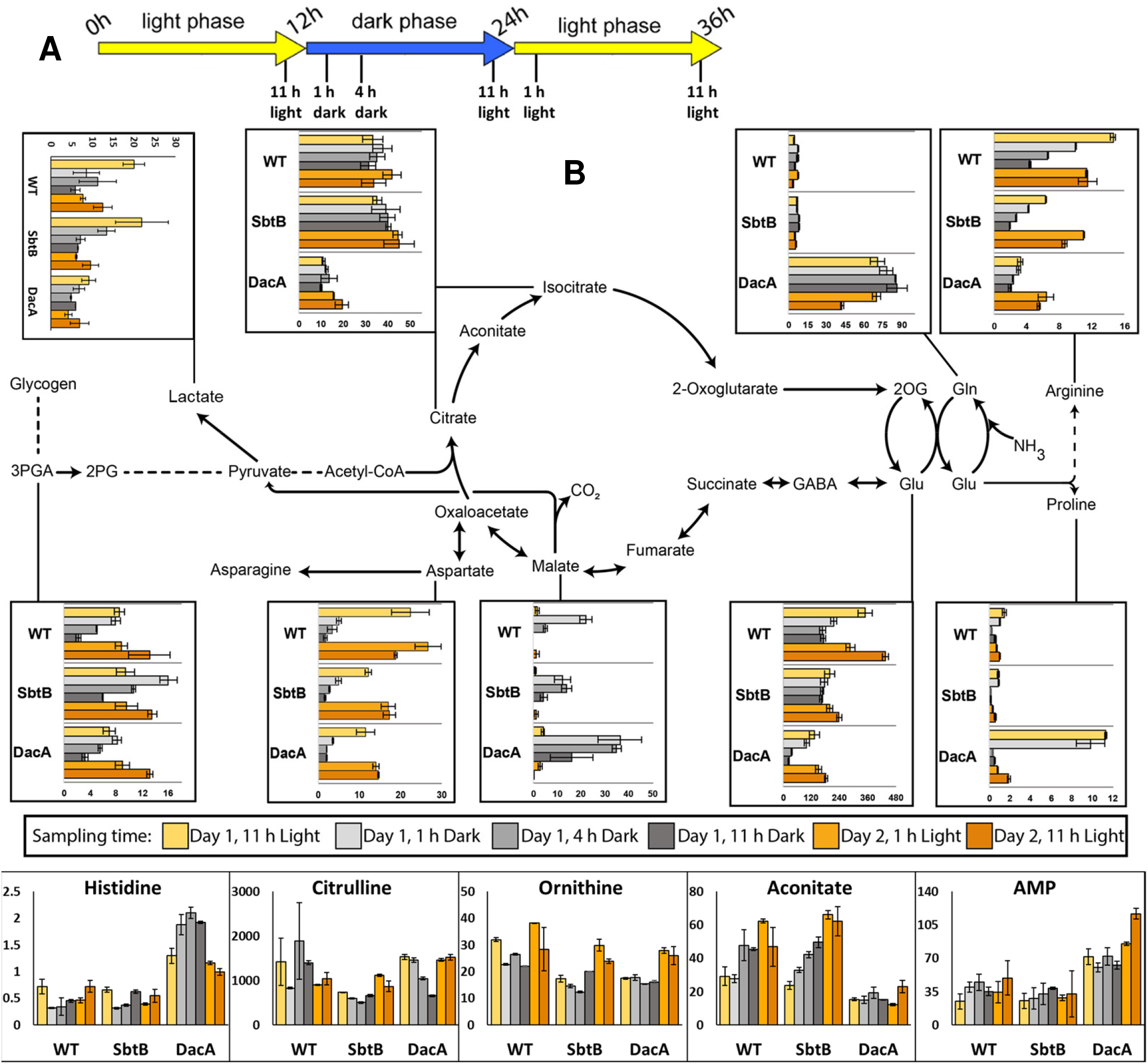
Targeted LC-MS based metabolomics analysis and additional metabolites of interest. **(A)** Schematic overview of the cultivation in the diurnal rhythm and sampling time points, as indicated. **(B)** Most significantly impacted metabolites of the central metabolism in WT, Δ*sbtB* and Δ*dacA* strains during 12 h day/12 h night diurnal rhythm, at 11 h of the first sampling day, 1, 4 and 11 h of the night-phase, and 1 and 11 h of the second sampling day. Metabolite levels are given as ng ml^-1^ OD750^-1^.

Interestingly, 3-phosphoglycerate (3PGA) levels did not change substantially between WT and mutants. This observation demonstrated that Ci acquisition is not the primary cause of the mutants’ growth defects. However, Δ*sbtB* other than WT cells did not decrease 3PGA content in the dark. This phenomenon can be explained on the bases of our recent finding that the diurnal switch of CCM activity is regulated via phytochromes involving SbtB (Oren et al. 2021). This explanation is also supported by our finding that SbtB controls a subset of LC-regulated genes and the bicarbonate flux in *Synechocystis* (Selim et al. 2018, Mantovani et al. 2022, Haffner et al. 2023). However, this could not exclude an alternation in the regulation of glyceraldehyde 3-phosphate dehydrogenase (GAPDH) and phosphoglycerate kinase or phosphoglycerate mutase (PGAM), the enzymes involved in producing and consuming 3PGA, respectively.

The most striking observation was concerning the glutamate (Glu) and glutamine (Gln) levels in both mutants (**Fig. 1, Supplemental Fig. S2 Table S2**). Decreased levels of glutamate in the light were already apparent in both mutants at the beginning of the experiment (11 h light of day 3). In the Δ*dacA* mutant, glutamate levels decreased to minimal levels by the end of the night phase (**Fig. 1**). On the other hand, this mutant accumulated 10-fold higher levels of glutamine both in the light and dark. This behavior was clearly specific for the Δ*dacA* mutant, since the Δ*sbtB* mutant showed glutamine balance similar to the wildtype (**Fig. 1, Supplemental Fig. S2)**. This result indicates a strong influence of c-di-AMP on nitrogen assimilation via the glutamine synthetase (GS) and glutamate synthase (GOGAT) cycle. The absence of c-di-AMP appears to dis-balance GS and GOGAT activities especially under dark conditions, at both the metabolome and transcriptome levels (Mantovani et al. 2022). The depletion of the Glu pool is consistent with the low osmotic stress tolerance of Δ*dacA* (Selim et al. 2021). The decreased Glu levels may be caused by decreased GOGAT turn-over or by increased anabolic consumption of Glu, since amino acids that require Glu for their biosynthesis (e.g. glutamine, histidine and proline) were significantly increased. Histidine accumulated predominantly during the night in Δ*dacA*. In contrast, proline levels in Δ*dacA* cells were significantly elevated at the end of day 3 and remained high during the first hour after dark shift (**Fig. 1**). Histidine depends on purine synthesis that requires glycine as one of the precursors, which was increased in the night similar to histidine in Δ*dacA* cells (**Fig. 1**). Low glutamate levels are known to stimulate the expression of proline synthesis genes and uptake. These mechanisms may contribute to high proline levels in Δ*dacA* (**Supplemental Fig. S2 and Table S2**). Proline is one of the most common compatible solutes, which accumulates as osmolytes under osmotic and salt stress, possibly as a compensatory effect of low Glu levels. Moreover, the metabolites associated with the ornithine–ammonia cycle (OAC), ornithine, citrulline, and arginine (**Fig. 1**) (Quintero et al. 2000, Zhang et al. 2018), that are derived from glutamate were partially altered, especially the arginine. Arginine levels were reduced in Δ*sbtB* cells and strongly depleted in darkened Δ*dacA* cells (**Fig 1**). This observation supported previously hypothesized arginine catabolism (Quintero et al. 2000) in the night and indicated that arginine utilization may be an early compensation of carbohydrate limitation (Selim et al. 2021) rather than a late, pleiotropic response of both mutants. The elevated levels of proline, which can be derived from arginine, could also explain the diminished arginine levels in Δ*dacA* mutant.

Targeted metabolomics revealed a unique difference of the Δ*dacA* as compared to Δ*sbtB* in regard to amino acid metabolism, especially decreased glutamate and increased glutamine, and in addition to the oxidative branch of TCA metabolism (**Supplemental Tables S1 and S2**). Intermediates of the oxidative branch, namely citrate/isocitrate and aconitate (**Fig. 1, Supplemental Fig. S2**), were clearly decreased in Δ*dacA* cells, which may explain a role of c-di-AMP in the supply of carbon precursors for glutamate synthesis. In contrast, the intermediates in the reductive branch of the TCA pathways appeared to accumulate in the mutant Δ*dacA* cells, as the intermediate malate was elevated in Δ*dacA* particularly during the night compared to WT and Δ*sbtB* cells (**Fig. 1**). Inversely to malate, aspartate and asparagine accumulated during the day and depleted towards the end of the night (**Fig. 1, Supplemental Fig. S1**). These trends were similar in both mutants and WT, but their levels were partially reduced in both mutants during the day. Aspartate is synthesized from oxaloacetate via transamination and Asn is synthesized form Asp by asparagine synthetase. Oxaloacetate can be produced by Ci-fixation via the phosphoenolpyruvate carboxylase reaction that may account for 20-25% of newly fixed Ci. This alternative Ci-fixation was apparently not strongly affected in both mutants. Interestingly, the Δ*dacA* cells weren’t able to activate the fermentative metabolism as evidenced by reduction of the fermentation product, lactate, compared to WT and Δ*sbtB* cells. Lactate production is accomplished with oxidizing NADH to NAD^+^. Hence, lower lactate in Δ*dacA* could indicate a redox imbalance with less excess of NADH for lactate fermentation. Also, we observed a strong accumulation of the aromatic amino acid, tyrosine, in Δ*dacA* mutant (**Supplemental Fig. S1)**.

Finally, we observed that AMP levels were significantly higher in the Δ*dacA* mutant (**Fig. 1**), further supporting the notion of energy depletion in Δ*dacA* cells, indicating a sort of global energy crisis. This could be due to several reasons, for instance, the impairment of glycogen anabolism and the elevated Gln biosynthesis, which depletes cellular ATP. Collectively, all these observations support the broader regulatory impact of c-di-AMP signaling in *Synechocystis* with dysregulation of various cellular metabolisms accompanied with c-di-AMP deficiency in Δ*dacA* cells, whereas SbtB seems more specific to central carbon metabolism.

### Proteomic landscape of Δs*btB* and Δ*dacA* mutants during the diurnal rhythm

To further elucidate how the deletion of SbtB or DacA causes the observed metabolic crisis in the respective mutants during diurnal growth, we performed a label-free quantitative proteome analysis. Cultures of the WT and both mutants were acclimated for two subsequent diurnal cycles and subsequently harvested at four time points (T1-T4) from day 3 covering one diurnal cycle, including midday (T1), midnight (T2), the end of the night (T3) and the following midday (T4) (**Fig. 2A**). For each strain, proteome analysis of three independent replicates were performed and displayed high reproducibility based on protein abundance correlations (**Supplemental Fig. S3**). Overall, 2,135 proteins could be identified in our proteome data set, of which 1,743 and 1,724 proteins revealed intensity ratios for at least one time point for Δ*sbtB*/WT and Δ*dacA*/WT, respectively (**Supplemental Table S3**). The overall protein abundances in the Δ*sbtB* mutant did not significantly differ from those of WT among all tested conditions (T1-T4), as indicated by co-clustering in a PCA (**Supplemental Fig. S4**). In contrast, protein abundances in the Δ*dacA* mutant were divergent from WT levels, indicating profound changes in the proteome as a potential result of reduced c-di-AMP availability (**Supplemental Fig. S4**). This is consistent to our targeted metabolomics analysis, indicating that the role of SbtB seems restricted to the central carbon metabolism, while a broad effect of c-di-AMP on both, the central carbon and nitrogen metabolism, became apparent.

**Fig. 2:**
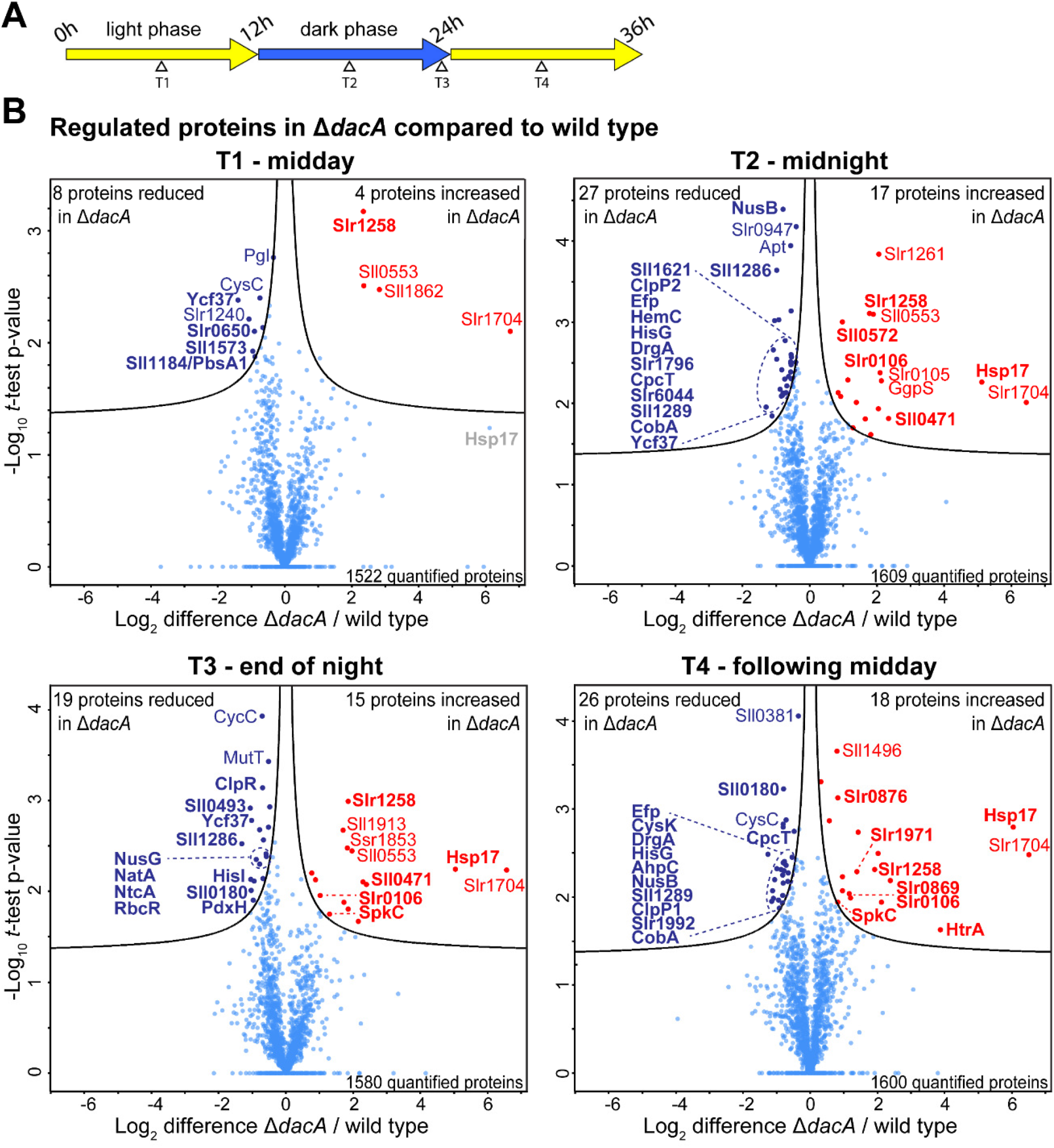
Proteome alterations of Δ*dacA* mutant. **(A)** Schematic overview of the cultivation in the diurnal rhythm and sampling at four different time points, with T1 representing midday, T2 midnight, T3 end of night and T4 the following midday. **(B)** Quantitative comparison of protein abundance between Δ*dacA* and wildtype at the four defined time points during the diurnal rhythm. Corresponding volcano plots indicate differences in protein abundance (Log_2_ difference Δ*dacA*/wildtype) and corresponding −log_10_ p-values from *t*-test of three independent replicates per strain and time point. Proteins with significant changes in abundance at an p-value threshold of 0.05 are labeled in blue (reduced in Δ*dacA*) and red (increased in Δ*dacA*).

To further specify proteins with significantly altered protein levels in Δ*dacA* or Δ*sbtB* relative to the WT, we performed student’s *t*-tests (p-value = 0.05) at all experimental conditions (**Supplemental Table S3**). In line with the PCA of the proteome, overall more proteins with significantly changed abundances could be detected in this analysis in Δ*dacA*, which in addition revealed higher-fold reduced or increased abundances compared to Δ*sbtB* (**Fig. 2, Supplemental Fig. S5**). To specify cellular processes that are potentially regulated by c-di-AMP, we grouped proteins in Δ*dacA* that showed a significant deviation from the WT as well as those proteins that followed the same trend in functional categories such as carbon and nitrogen metabolisms, ion transport and homeostasis, photosystem assembly, redox regulation, as well as transcriptional regulation (**Supplemental Fig. S6A-E and Supplemental Table S3**). Thereby, regulated proteins reported as “hypothetical” or “unknown” were blasted against the UniProt Reference Clusters (UniRef) and the RCSB Protein Data Bank (PDB) to retrieve potential cellular functions based on sequence and structural homologies.

Consistent with the metabolome analysis, which showed impaired regulation of the GS-GOGAT cycle metabolites, proteomic levels of the Δ*dacA* mutant showed striking changes in nitrogen metabolism related proteins (**Fig 2, Supplemental Fig S6A**). These changes were marked by a reduced abundance of the cysteine synthetase CysK (*slr1842*), the porphobilinogen deaminase HemC (*slr1887*), the nitrogen utilization protein NusB (*sll0271*), the histidine biosynthesis enzymes HisG (*sll0900*) and HisIE (*slr0608*), as well as the NrtC subunit of nitrate/nitrite transporter (*sll1452*), and NatA (*slr0467*) and LivF (*slr1881*) subunits of amino acids transporters **(Supplemental Table S3)**. In addition, the protein levels of the key enzyme of arginine biosynthesis N-acetyl-L-glutamate kinase (NAGK; *slr1898*; Lapina et al. 2018, Selim et al. 2019, 2020a & 2020b) were slightly reduced in Δ*dacA*, which could explain the reduction of arginine levels in the Δ*dacA* mutant. Interestingly, also the global nitrogen regulator NtcA (*sll1423*), the master transcription factor of nitrogen metabolism (Forchhammer and Selim 2020, Selim et al. 2019), was reduced in Δ*dacA* among all conditions down to 1.73-fold in the end of the night phase (**Fig 2, Supplemental Fig. S6A**). During nitrogen depletion, NtcA in complex with its coactivator PipX induces genes required for nitrogen assimilation (Selim et al. 2019, LIop et al. 2023), such as glutamine synthetase (GS; *slr1756*), ammonium transporter (AMT1; *sll0108*) and urea transporter, and represses genes that inhibit N-assimilation, such as the GS inhibition factor 17 (IF17; *sll1515*) and PirA (*ssr0692*) (Giner-Lamia et al. 2017). PirA is known to regulate the flux into the ornithine-ammonia cycle (Bolay et al. 2021). Under nitrogen excess, however, free NtcA does not activate the expression of nitrogen metabolism related genes (Fadi et al. 2003, Luque et al. 1994). As the GS synthesizes Gln from Glu and ammonia, the imbalance in the intracellular Gln/Glu ratio, comprising high Gln and low Glu levels, is consistent with the partial elevation of GS and AMT1 levels in Δ*dacA* cells (**Supplemental Table S3**). Concomitantly to the reduced levels of NtcA in Δ*dacA*, protein levels of both PirA and IF17 were increased (**Supplemental Fig. S6F**). The elevated PirA levels is in agreement with our transcriptome analysis revealing a low expression of the NtcA-dependent sRNA, NsiR4, the repressor of *pirA* (Bolay et al. 2022), which further explains the low Arg levels in Δ*dacA*. The induction of IF17 is consistent with the increased levels of glutamine, since expression of *gifB* (encoding IF17) is mediated by a glutamine-dependent riboswitch (Klähn et al. 2018). Thus, the elevation of IF17 represents a compensatory mechanism to prevent extra-flow of nitrogen into Gln by the activity of GS. Lower levels of the urea transporter subunit UrtA (*slr0447*) are also consistent with the decreased NtcA levels and the high nitrogen-flow, indicated by elevated Gln levels (**Fig. 1**). Also, we observed dysregulation of other proteins which are under the control of NtcA, such as the substrate binding subunit of polar amino acids transporter (*sll1762*) and hypothetical Slr1852 protein were reduced in abundance, while the anti-sigma F factor antagonist (*slr1912*) and the NblD (known also as Nsir6) revealed increased abundances (Baumgartner et al. 2016, Giner-Lamia et al. 2017, Krauspe et al. 2021). Altogether, the dysregulation of NtcA regulated genes in Δ*dacA* implies that Δ*dacA* is unable to sense and integrate correctly the nitrogen levels inside the cells and explains the impact of c-di-AMP on the central nitrogen metabolism in *Synechocystis*, including GS activity as well as the regulation of ornithine-ammonia cycle. In addition, it hints at an additional regulatory input in NtcA-mediated gene expression that is affected by c-di-AMP.

In the second group, several uncharacterized proteins with high similarities to membrane associated transporters were grouped. Among these is a potential member of the dynamin family (*slr0869*) and a hypothetical protein (*slr1258*), which contains a SIMPL-like domain, both with higher abundance levels in Δ*dacA* as well as a potential efflux transporter (*sll0180*) with overall reduced levels. Interestingly, the hypothetical proteins Slr6044 (*slr6044*) and Sll0493 (*sll0493*) revealed high similarities to the copper transporter subunit CopC and the sodium dependent K^+^-transporter subunit KtrA, respectively. These proteins were significantly reduced protein levels in Δ*dacA* (**Fig. 2, Supplemental Fig. S4B**). Remarkably, the KtrA was previously described as a c-di-AMP target protein in *Synechocystis* (Selim et al. 2021). Interestingly, the glucosylglycerol-phosphate synthase (GgpS; *sll1566*), a key enzyme for synthesis of the dominate compatible solute glucosylglycerol in *Synechocystis* and many other cyanobacteria (Kirsch et al. 2019), is found at elevated levels in mutant Δ*dacA* cells. This is in strong agreement with previous reports of the role of c-di-AMP in ion homeostasis and osmoprotection in cyanobacteria (Selim et al. 2021, Agostoni et al. 2018).

Furthermore, we found proteins with significantly changed abundances to be involved in photosystem assembly and maintenance of photosynthesis (**Fig. 2, Supplemental Fig S4C**). Among this functional group, proteins involved in photosystem I (PSI) assembly are differentially regulated while genes encoding proteins related to photosystem II remained largely unaffected. For example, the green-light chromophore domain containing linker protein of PSI antenna complex CpcL (*cpcG2*; *sll1471*) (Kondo et al. 2007) and the PSI assembly related proteins Ycf37 (*slr0171*) (Wilde et al. 2001) were strongly reduced up to 1.74- and 2.65-fold in Δ*dacA*, respectively. Moreover, other proteins related to the assembly of photosystems, namely Ycf54 (*slr1780*), Ycf53 (*sll0558*), Ycf3 (*slr0823*), Ycf58 (*slr2049*), Sll0226 protein, PsaL (*slr1655*), PSI biogenesis protein BtpA (*sll0634*), phycobilisome rod linker CpcC2 (*sll1579*) and the response regulator for the energy transfer from phycobilisomes to PS (*slr0947*), also were dysregulated relative to the WT cells (**Supplemental Table S3**). In addition, the hypothetical proteins Slr1649 and Sll1573, which are related to the chromophore lyase CpcT/CpeT and the chromophore lyase CpcS/CpeS, consecutively, were reduced in Δ*dacA*. The impairment in PSI assembly and photosynthesis, caused by lower amounts of PSI biogenesis and assembly proteins upon c-di-AMP depletion, hints at imbalance in cyclic electron flow, which might also contribute and explain the impairment of Δ*dacA* under diurnal growth and the reduction of photosystem quantum yield (Selim et al. 2021), and thereby the observed energy crisis as indicated by the elevation of AMP levels (**Fig. 1**).

The deficiency in photosystem biogenesis is accompanied by a reduction of several proteins, involved in metabolism of cofactors and redox regulation (**Fig. 2, Supplemental Fig. S4D**), such as proteins of the thiol-specific antioxidant AhpC/TSA family (*sll1289*, *sll1621*), thioredoxin Slr1796 (*slr1796*), the heme oxygenase PbsA1 (*sll1184*), the pyridoxamine-5-phosphate oxidase PdxH (*sll1440*), and the glutathione peroxidase like NADH peroxidase Gpx2 (*slr1992*). The cob(I)alamin adenosyltransferase CobA (*slr0260*), IsiB flavodoxin (*sll0248*), and the DrgA nitro/oxidoreductase (*slr1719*) followed also that trend (**Supplemental Table S3**). However, the hypothetical proteins of oxidoreductase family (*sll0471* and *slr0876*), were increased in the Δ*dacA* mutant (**Supplemental Table S3**). In accordance with those changes, we detected an up to 27-35% increase in the production of reactive oxygen species (ROS) during the day and night phases, respectively, for both Δ*dacA* and Δ*sbtB* mutants relative to WT cells (**Fig. 3**). The increase in ROS production serves as a delicate explanation for the observed metabolic increase of glutathione (**Fig. 3**), which functions as a cellular response to elevated intracellular redox stress (Tel-Or et al. 1985, Li et al. 2007, Latifi et al. 2009). The Δ*dacA* and Δ*sbtB* cells showed almost 3-fold and 2-fold increase in the glutathione levels relative to wildtype, respectively (**Fig. 3**). Furthermore, elevated ROS and glutathione levels is another aspect of the metabolic crisis, which the mutants undergo in diurnal growth and which, besides others, ultimately leads to cell death, similar to what observed in the mutant of the circadian clock regulator, RpaA (Diamond et al. 2017, Scheurer et al. 2021). Notably, our proteomics dataset revealed a slight reduction of RpaA (*slr0115*) levels in Δ*dacA* (**Supplemental Table S3**).

**Fig. 3:**
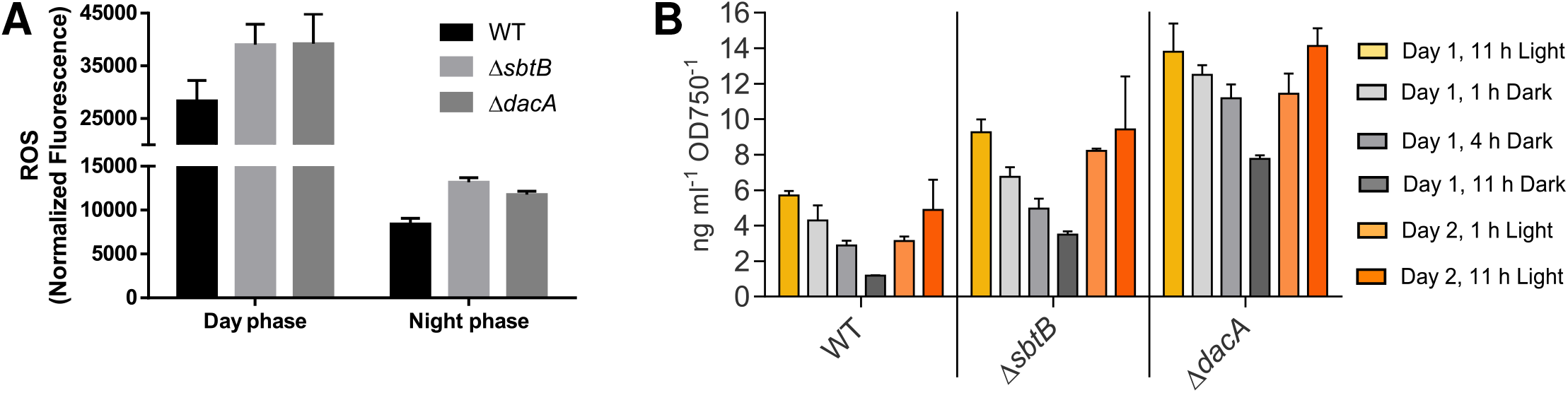
Reactive Oxygen Species (ROS) and Glutathione quantifications. **(A)** Intracellular ROS production of *Synechocystis* WT, Δ*sbtB*, and Δ*dacA* during the day and night phases of diurnal growth, showing high levels of ROS during the day phase due to the photosynthetic activity with an increase of ≈ 30% increase in both mutants. In the night phase, the levels of ROS drop, but both mutants showed still an elevation by ≈ 30% in ROS compared to WT cells. **(B)** Intracellular glutathione levels of *Synechocystis* WT, Δ*sbtB*, and Δ*dacA* during diurnal cycles, as indicated and showing an elevated levels of glutathione in both mutants, but especially in Δ*dacA* compared to WT cells.

The broad range of cellular processes affected by c-di-AMP is also reflected by various proteins involved in transcription, translational and posttranslational regulations, which were dysregulated in Δ*dacA* (**Fig. 2, Supplemental Fig. S4E**). Among those proteins, the transcriptional termination factor NusG (*sll1742*), the elongation factor P (*slr0434*), the transcriptional regulators (*sll1286* and *sll0998*) with high similarity to TetR and LysR, respectively, as well as the hypothetical protein Slr0650, which contains a NYN-domain, were reduced, while the Rho termination factor (*slr0106*), the potential GTPase Sll0572 (*sll0572*) and the serine/threonine kinase SpkC (*slr0599*) were significantly increased in Δ*dacA*. As indication of proteome instability in Δ*dacA*, we also observed a strong increase in the stress-response chaperone Hsp17 (*sll1514*), the peptidase Slr1971, and the HtrA protease (*slr1204*), while the Clp proteases (*slr0542*, *sll0534*, *slr0164*, *slr0165*) and the periplasmic YmxG protease (*slr1331*) were reduced (**Supplemental Table S3**). In *Synechocystis*, the Hsp17 chaperone is known to stabilize the stressed membranes and to bind the denatured proteins for subsequent refolding (Török et al. 2001), consistent with the strong increase of HtrA protease. The HtrA proteases are generally implicated in stress responses, oxidative damage and photosynthesis (Clausen et al. 2011, Huesgen et al. 2011), supporting the notion of a strong cell stress as a result of e.g. oxidative damage (**Fig. 3**), and further explains the lethality associated with *dacA* mutation. Although the exact consequence of these alterations in the transcriptional and translational machineries are unclear, their existence indicates a pleiotropic effect of c-di-AMP on cellular processes, which needs to be deciphered.

### Physiological relevance of c-di-AMP for nitrogen regulation

To further validate the physiological relevance of c-di-AMP for nitrogen metabolism, we estimated cellular levels of cyanophycin, a nitrogen-rich reserve biopolymer synthesized from arginine and aspartate, in Δ*dacA* and WT. Cyanophycin is also considered as a stress marker in cyanobacteria as it accumulates under various stress conditions of nitrogen, potassium and phosphate starvations, salt stress, and diurnal growth (Page-Sharp et al. 1998, Watzer et al. 2015, Watzer and Forchhammer 2018). The intracellular cyanophycin levels were elevated in Δ*dacA* after 9 days of diurnal growth compared to WT and Δ*sbtB* cells, however we were not able to confirm that microscopically using Sakaguchi staining (**Fig. 4A,B**). To support the tendency of more cyanophycin in Δ*dacA*, we combined the diurnal growth with phosphate starvation, a condition that favors cyanophycin accumulation (Stevens et al. 1981, Trautmann et al. 2016). As expected the amount of cyanophycin was significantly increased in Δ*dacA* over WT and Δ*sbtB* cells (**Fig. 4C,D**). The fact, that cyanophycin levels were increased in Δ*dacA* cells serves as additional evidence for the role of c-di-AMP in *Synechocystis* nitrogen metabolism and further support the stressed state of Δ*dacA* cells. The higher accumulation of cyanophycin could drain and thereby decrease the Arg levels in Δ*dacA* as observed in our metabolome analysis.

**Fig. 4:**
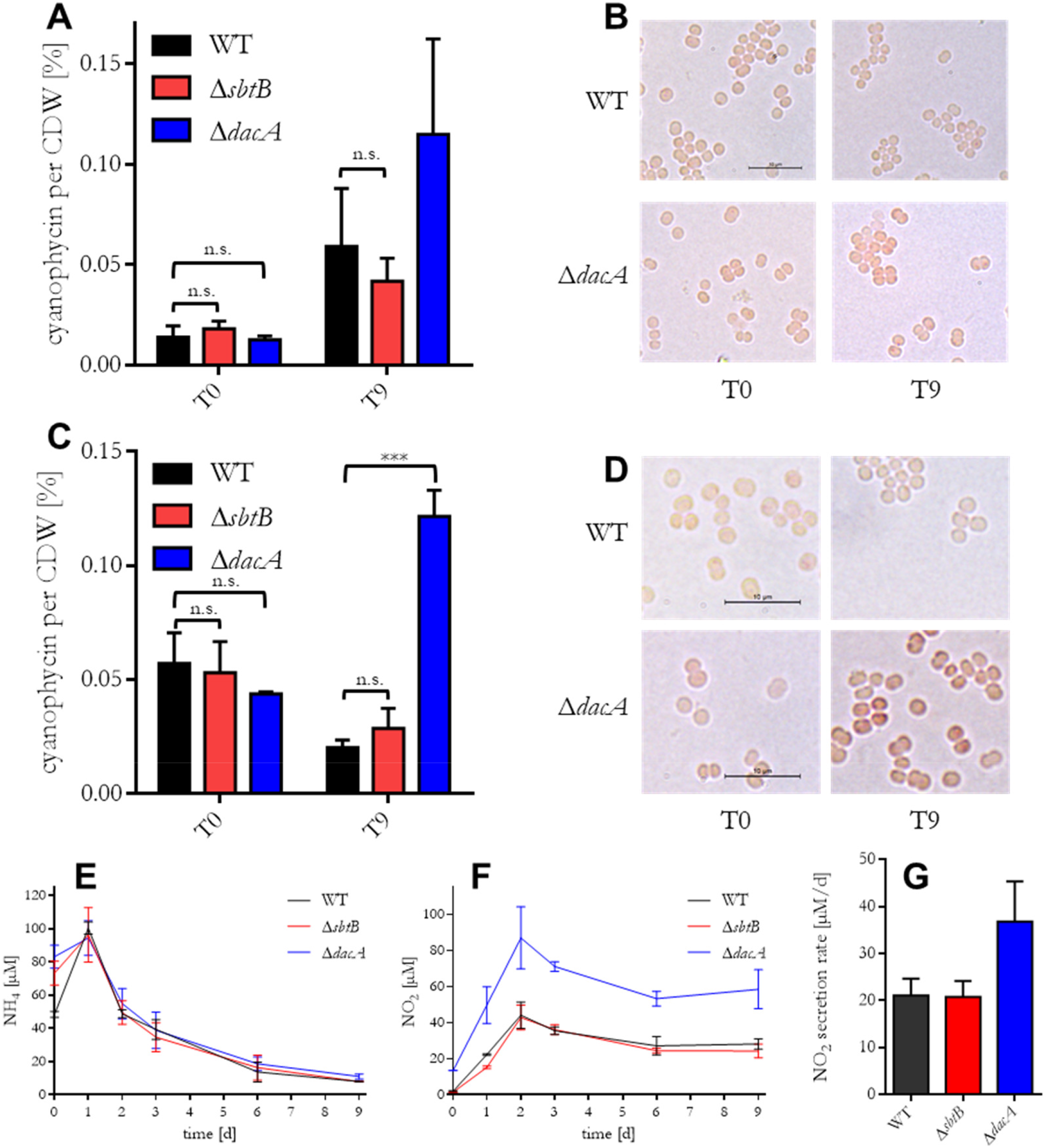
Physiological relevance of c-diAMP for nitrogen metabolism. **(A, C)** Cyanophycin quantification of Δ*sbtB* (red) and Δ*dacA* (blue) in comaprison to *Synechocystis* WT before (T0; left pannel) and 9 days after the shift to diurnal growth without **(A)** and with additional phosphat starvation **(C)** (T9; right pannel). **(B, D)** Light microscopic pictures of sakaguchi stained cyanophycin granules (resulting in intracellular red spots) of *Synechocystis* WT (top pannel) and Δ*dacA* (bottom pannel), before (T0; left pannel) and 9 days after the shift to diurnal growth without **(B)** and with additional phosphat starvation **(D)** (T9; right pannel); scale bar: 10 µM. **(E, F)** Ammonium [NH_4_^+^] **(E)** and nitrite [NO_2_^-^] **(F)** quantification (in [µM]; y-axis) of Δ*dacA* (blue) and Δ*sbtB* (red) in comparison to *Synechocystis* WT (black) throughout 9 days of diurnal growth (x-axis). **(G)** NO_2_^-^ secretion rate (in [µM/d]; y-axis) of of Δ*dacA* (blue) and Δ*sbtB* (red) in comparison to *Synechocystis* WT (black) throughout the first two days of diurnal growth, based on the data shown in (F).

The proteome analysis revealed also changes in transporter proteins related to nitrogen and amino acid uptake. Therefore, we analyzed to which extent c-di-AMP is involved in nitrogen uptake and reduction. To this end, we measured extracellular ammonium (NH_4_^+^) and nitrite (NO_2_^-^) levels of Δ*dacA*, Δ*sbtB*, and WT throughout 9 days of diurnal growth during the day phases (**Fig. 4E-G**). Ammonium levels of WT peaked in the first day of diurnal growth and then steadily decreased over the following 8 days. An identical pattern was found in the mutants (**Fig. 4E**). In contrast to ammonium, a significant increase of extracellular NO_2_^-^ levels were observed in Δ*dacA* as compared to WT and Δ*sbtB*. After two days of diurnal growth, nitrite levels of WT and Δ*sbtB* reached their plateau at 44.1 ± 7.3 µM and 42.9 ± 6.9 µM, respectively, while the plateau value for Δ*dacA* was nearly two-times higher and reached at 87.0 ± 17.2 µM (**Fig. 4F**). The respective NO_2_^-^ secretion rate throughout the first two days of diurnal growth of Δ*dacA* was 75% higher than those of WT and Δ*sbtB* (**Fig. 4G**). For nitrogen assimilation, NO_3_^-^, the main inorganic nitrogen source in the BG11 medium, is taken up by *Synechocystis* via the NrtABCD transport system (Omata 1995, Koropatkin et al. 2006) and gets reduced to NO_2_^-^ and ultimately to ammonia (NH_3_), which is finally assimilated in the GS/GOGAT cycle (Bolay et al. 2018). Nitrite excretion can be a consequence of overactive nitrate uptake and reduction relative to nitrite-reductase activity (Kloft and Forchhammer 2005), which could further influencing on GS-GOGAT cycle and thereby on the overall Gln/Glu levels (**Fig. 1**). Additionally, the toxicity of NO_2_^-^ towards *Synechocystis* serves as another explanation for Δ*dacA* lethality (Selim et al. 2021), as it was reported that elevated levels of nitrite is influencing negatively on photosystem II (PSII) activity (Zhang, Ma et al. 2017), similar to what we observed for Δ*dacA* with 25% reduction of PSII quantum yield (Selim et al. 2021).

### SbtB interactome extends to further bicarbonate transport systems

While both of our metabolomics as well as proteomics analysis indicates a broad effect of c-di-AMP on a great variety of cellular processes, the function of SbtB seems to be restricted to the central carbon metabolism. Accordingly, we detected an increase of CmpA (*slr0040*) levels in Δ*sbtB* in comparison to WT in our proteomics analysis (**Supplemental Fig. S7A**). CmpA is the substrate binding subunit of the ABC-type bicarbonate transporter complex BCT1, which is encoded by the *cmpABCD* operon (*slr0040*-*slr0044*) (Koropatkin et al. 2007). Interestingly, the CmpABCD complex belongs to the same protein family as the nitrate/nitrite ABC-type transporter NrtABCD (Koropatkin et al. 2006), which was shown to be regulated by canonical PII via physical interaction with NrtC/D subunits (Watzer et al. 2019). Due to high structural similarity between SbtB and canonical PII proteins, binding of the PII-like protein SbtB to the CmpABCD complex seemed, therefore, likely. To test our hypothesis, we performed a Ni^2+^-NTA based pulldown analysis of immobilized C-terminal His_8_-tagged SbtB with *Synechocystis* crude cell extracts, either in the presence or absence of 100 µM c-di-AMP. Another C-terminal His_8_-tagged protein, Slr1970, was used as a negative control. Indeed, these pulldowns revealed the BCT1 subunits CmpA, CmpC, and CmpD as potential SbtB-interaction partners. Surprisingly, the interaction of SbtB with CmpA, CmpC and CmpD was enriched in the presence of c-di-AMP (**Supplemental Fig. S7B,C**). This agrees with the previous observation that the expression of both the *cmp* operon was dysregulated in Δ*dacA* (Mantovani et al. 2022). To confirm the result of the pulldown, we performed bacterial two hybrid (B2H) assays (**Supplemental Fig. S7D**). The B2H assays revealed interaction of C- and N-terminal tagged SbtB (SbtB-T25 and T25-SbtB, respectively) fusion proteins with either C- or N-terminal tagged CmpC (CmpC-T18, T18-CmpC) and C-terminal tagged CmpD (CmpD-T18). A strong interaction of SbtB with CmpC was only detected between N-terminal tagged T25-SbtB with C-terminal tagged CmpC-T18, while a strong interaction of CmpD-T18 was only visible upon co-expression with C-terminal SbtB-T25 (**Supplemental Fig. S7D)**. The interaction of SbtB with the cytosolic subunits of the BCT1 complex shows that the role of SbtB in bicarbonate uptake is not solely restricted to the regulation of SbtA (Selim, et al. 2023, Haffner et al. 2023).

The cyanobacterial CCM includes a third bicarbonate uptake system, BicA, which also shows a different expression pattern in the Δ*sbtB* knockout mutant (Selim et al. 2018). However, BicA didn’t pulldown with the SbtB. BicA is a member of SLC26 family of anion transporters with a large cytoplasmic STAS-domain (antisigma factor antagonist), required for regulation of the transport activity via interaction with other regulatory proteins (Babu et al. 2010, Geertsma et al. 2015, Wang et al. 2019). Therefore, we checked additionally for a possible interaction of SbtB with BicA by B2H assays. In these B2H assays, a strong positive signal was observed for C-terminal tagged BicA-T18 with either C- or N-terminal tagged SbtB-T25 (**Supplemental Fig. S7D)**. Collectively, these analyses revealed that SbtB is not only restricted to the regulation of SbtA but has a broader role in controlling the major HCO_3_^-^ transporters of the cyanobacterial CCM (Selim et al. 2018, Mantovani et al. 2022 & 2023a). However, the molecular details of these interactions and the influence of SbtB on BCT1 and BicA activity need further examination.

## Remarks & conclusions

Our metabolome characterization of the c-di-AMP synthase mutant Δ*dacA* and the c-di-AMP receptor mutant Δ*sbtB* points to a pronounced imbalance of carbon/nitrogen homeostasis in the absence of the signaling molecule c-di-AMP (**Fig. 5**). This imbalance progresses under light-dark cycles and appears to cause common compensation responses. The metabolome of Δ*dacA* cells is considerably different from the Δ*sbtB* mutant already at early stages after a shift to light-dark cycles that are ultimately detrimental to both mutants. These observations point to a broader impact of c-di-AMP deficiency than mediated by the lack of SbtB signaling alone and reinforces our previous results that c-di-AMP targets more metabolic regulators than just SbtB (**Fig. 5**) (Selim et al. 2021). The deficiency of glycogen accumulation caused by deletion of the c-di-AMP receptor SbtB appears to drive late pleiotropic metabolic responses in both mutants, causing growth impairment under diurnal rhythm. In general, marked changes were observed mainly to metabolites related to TCA cycle, amino acids, nitrogen metabolism, and photorespiration (**Fig. 1**), implying a broad impact of Δ*dacA* mutation and to some extent Δ*sbtB*, on central carbon and nitrogen metabolisms than only on glycogen metabolism or as a consequence of lower glycogen levels in the mutants (**Fig. 5**). The most striking differences shared between both mutants in comparison to WT cells were related to amino acids (**Fig. 1**), such as the reduction of glutamate, a key amino acid in nitrogen metabolism. We interpret these observations as a penultimate mechanism that mobilized carbon from amino acid bound resources to counterbalance long-term and aggravating carbohydrate limitation of both mutants when shifted from constant light to day-night cycles.

**Fig. 5:**
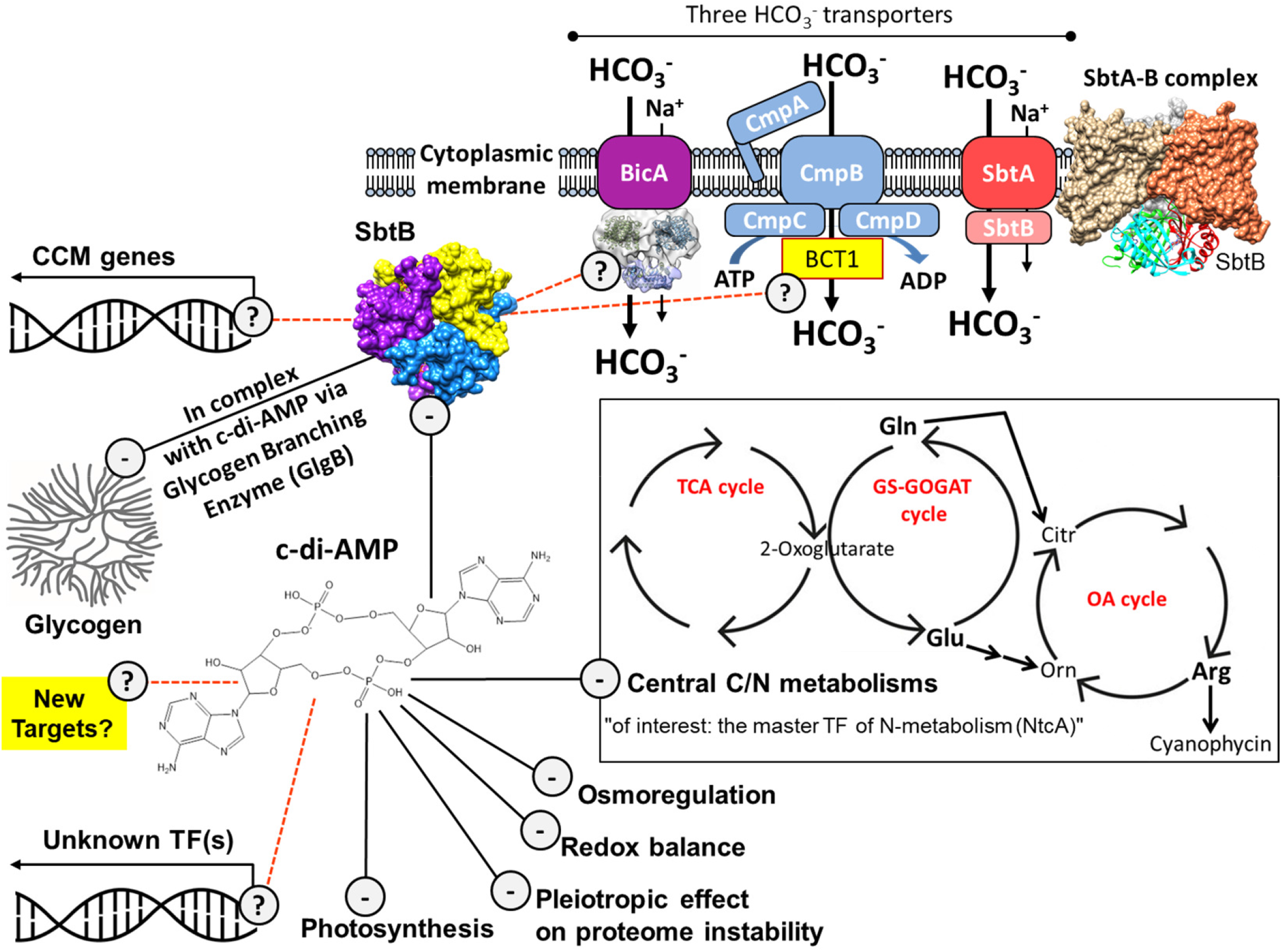
Model for regulation of cyanobacterial central metabolism by c-di-AMP and SbtB signaling in response to diurnal rhythms. The negative consequences of the absence of c-di-AMP in Δ*dacA* mutant on various cellular processes is indicated by (-). Of particular importance the influences of c-di-AMP on central C/N metabolisms of TCA, GS-COGAT and ornithine–ammonia (OA) cycles. The possible cellular effects of c-di-AMP are likely to be mediated through the binding to yet undiscovered transcription factor(s) (TF) or other new targets, as indicated by (?). On the other hand, SbtB seems to work specifically, but broadly, on central carbon metabolism by regulating the glycogen biosynthesis and the major bicarbonate transporters (SbtA, BicA and CmpABCB complex) through direct protein interaction (Selim et al. 2018 & 2021, Haffner et al. 2023). However, the exact regulatory modes of SbtB on BicA and BCT1 complex remain elusive, as indicated by (?). SbtB regulates glycogen anabolism through the interaction with the glycogen branching enzyme (GlgB) in a c-di-AMP-dependent manner, therefore the absence of either SbtB or c-di-AMP influences negatively on glycogen synthesis, indicated by by (-). SbtB also influences on C-homeostasis by controlling the CCM genes expression (Mantovani et al. 2022), possibly through an interaction with yet unidentified transcription factor(s), indicated by (?).

Further, we confirmed the essential role of c-di-AMP for several cellular processes such as ion homeostasis, redox homeostasis, transcription, and transitional regulations, photosynthesis, and especially in central nitrogen metabolism by differential proteome characterization (**Figs. 2 & 5**). These combined analyses permitted to link the intracellular glutamine/glutamate imbalance, reduced arginine in Δ*dacA* mutant with the decreased expression of the global nitrogen transcription factor NtcA. However, based on our data sets, we could not clarify, whether the cellular levels of NtcA is directly linked to c-di-AMP or if it is a downstream effect of the carbon metabolism dysregulations. But, our results suggested an additional regulatory input in NtcA-mediated gene expression that is affected by c-di-AMP, despite, we were able to exclude a direct binding of c-di-AMP to NtcA. Nevertheless, our data revealed a broad range of cellular processes that are subjected to c-di-AMP regulation, likely through binding to yet unidentified transcription factor(s) or small RNAs to propagate the c-di-AMP-mediated responses. In sum all these processes might lead to the observed lethality of the c-di-AMP deficient Δ*dacA* mutant under diurnal growth, while the diurnal lethality of Δ*sbtB* seems to be restricted to carbon metabolism (Selim et al. 2021, Mantovani et al. 2022 & 2023a).

## Materials and Methods

### Cultivation conditions

Cyanobacterial growth experiments were performed as previously described in (Selim et al. 2018, 2021 & 2023). Cells were always normalized to their optical density at 750 nm using a Helios Gamma UV-Vis Spectrophotometer (Thermo Fisher Scientific). Experiments in day-night conditions were performed in a separate day-night chamber, providing 12-hour light phase, followed by a 12-hour darkness phase.

### Diurnal rhythm metabolome (Targeted metabolomics) analysis

The study of the metabolome during diurnal rhythm was started with the pre-culturing of the strains in a Multicultivator MC-1000 (PSI, Czech Republic) with a 12 h light (130 µmol photons m-2 s-1)/12 h dark cycle with an OD_720_=1 under low carbon (LC; ambient air) conditions. After acclimation to diurnal rhythm for 2 days, cultures were normalized to an OD_750_=1 in bubbling tubes with a 12 h day/12 h night rhythm. Samples were taken at 11 h of the first photoperiod, then 1, 4 and 11 h of the dark period, and finally 1 and 11 h of the second photoperiod. Sampling was performed via vacuum filtration on nitrocellulose filters with a pore size of 0.45 µm (Mackerey-Nagel, Germany), which were frozen in liquid nitrogen and stored at −80 °C.

For metabolites extractions, 730 µL LC-MS grade methanol with 1 µL of carnitine (1 mg/mL) were added to each filter, then shaken for 15 min at room temperature (RT). 400 µL chloroform was then added, followed by incubation at 37 °C for 10 min and further addition of 800 µL of LC-MS grade H_2_O. After shortly shaking via vortex, the samples were incubated over night at −20°C. The following day each sample was centrifuged at 14.000 rpm for 5 min at 4 °C. The upper phase was transferred in 1.5 mL tubes and the samples dried via overnight (ON) incubation in a Speed-Vac (Eppendorf, Germany). The samples were then re-suspended in 600 µL LC-MS grade H_2_O and filtered through 0.45 µm cut-off polyamide syringe filters (Mackerey-Nagel, Germany). The filtered supernatants were analyzed via LC-MS as previously described (Mantovani et al. 2022).

### Proteomic analysis

Cells of the wild type strain and the mutants were grown for proteomics analyses in 250 mL batch cultures under constant light until OD_750_= 0.25 and subsequently acclimated to diurnal rhythm for 2 days as described above. The cells were harvested by centrifugation (4.000 × g, 4°C, 10 min) from day 3 at midday (T1), midnight (T2), the end of the night (T3) and the following midday (T4) of day 4. Cell pellets were snap frozen and stored at −80°C, until proteins were extracted in 500 µL lysis buffer, reduced and alkylated, before proteins were precipitation with acetone/methanol as described before (Spät et al. 2015). Purified proteins were re-dissolved in denaturation buffer (6 M urea, 2 M thiourea in 100 mM Tris/HCl, pH 8) and concentrations were measured by Bradford assay. 20 µg of protein were separated and pre-digested with 200 ng Lys-C (Waco) for 3 h, then 200 ng trypsin (Promega), diluted in 80 µL of 20 mM ammonium bicarbonate buffer; pH 8.0, was added. Resulting peptide solutions were acidified with trifluoroacetic acid to pH 2.5 and de-salted via Stage tips (Ishihama, Rappsilber et al. 2006).

For each sample, 250 ng of the peptide mixture was subjected to LC-MS/MS analysis in a randomized order. each sample was loaded onto an in-house manufactured nanoLC column (20 cm; 70 µm ID; packed with ReproSil-Pur C18 1.9 µm particles; Dr. Maisch, Germany) and separated by RP-chromatography on an EASY-nLC 1200 (Thermo Fisher Scientific, USA) using 90 min linear gradients. Eluting peptides were on-line ionized in an ESI source and analyzed in the data-dependent acquisition mode on an Orbitrap Exploris 480 mass spectrometer (Thermo Fisher Scientific, USA) as described before (Spät et al. 2021).

All raw spectra were processed with the MaxQuant software (version 1.6.8.0) at default settings and enabled match between run option and label-free quantification (LFQ). Peak lists were searched against a target-decoy database of *Synechocystis* sp. PCC 6803 including 3680 protein sequences from Cyanobase and recent studies (Mitschke et al. 2011, Kopf et al. 2014, Baumgartner et al. 2016). False discovery rates were limited to 1% on peptide and protein levels. Perseus software suite (version 1.6.5.0) was used for statistical analyses of the LFQ dataset. Student’s *t*-tests for determination of significantly regulated proteins was performed at a p-value threshold for 0.05 with S_0_ of 0.5.

### Ammonium, nitrite, and nitrate quantification

To estimate the composition of extracellular combined nitrogen sources, 2 mL of *Synechocystis* culture were harvested for each time point via centrifugation at 4.000 g and 4 °C for 10 min. Clear supernatants of each culture were used for ammonium quantification via Nessler reaction (Vogel et al. 1989), as well as nitrite quantification via Griss reaction (Fiddler 1977) and nitrate quantification via estimation of absorbance at 210 nm (Kloft and Forchhammer 2005).

### Cyanophycin extraction and quantification

Cyanophycin extraction and quantification was performed as previously described in (Elbahloul et al. 2005). Cell pellets gathered from 50 mL culture volume were resuspended in 1 mL acetone and incubated for 30 min at 1.400 rpm shaking. After 10 min of centrifugation at 13.000 rpm pellets were resuspended in 1.2 mL of 100 mM HCl and incubated for 2 h at 40 °C. After 10 min of centrifugation at 13.000 rpm and 4 °C, pellets were discarded and 300 µL of 1 M Tris/HCl pH 8.0 were added to supernatants, followed by an overnight incubation at 4 °C. Precipitated cyanophycin was pelleted at 18.000 rpm for 15 min. Cyanophycin quantification was performed via Bradford reagent at 595 nm and normalized to cell dry weight.

### Cyanophycin staining using the Sakaguchi reaction

Intracellular cyanophycin was stained via the Sakaguchi reaction according to (Messineo 1966). Therefore, cell pellets of 500 µL culture were fixated in 500 µL PBS [2,7mM KCl; 1,5mM KH2PO4; 137mM NaCl; 8,1mM Na2HPO4; pH 7.4] with additional 2.5% (w/v) glutaraldehyde for 30 min on ice. The fixation was stopped by pelleting the cells at 8.000 g and 4 °C for 2 min and washing the pellets one time with 500 µL PBS. To perform the Sakaguchi reaction, fixated cells were centrifuged at 8.000 g for 2 min and pellets were resuspended in 80 µL of 5 M KOH. After adding 20 µL of 1% (w/v) 2,4-Dichloro-1-naphthol solution, cells were incubated for 5 min at room temperature. Immediately after the addition of 20 µL 5% (v/v) sodium hypochlorite, cells were centrifuged at 4.000 g for 5 min. Stained cell pellets were resuspended in 20 µL PBS whereof 5 µL were used for light microscopy on 2% agarose covered blades.

### Quantification of Reactive oxygen species (ROS)

Reactive oxygen species were quantified as previously described in (Diamond et al. 2017), using the H2DCFDA fluorescent marker. H2DCFDAwas added to 1 mL *Synechocystis* culture to a final concentration of 5 µM and incubated for 30 min at 30 °C, protected from light. 200 µL of each sample were transferred into a 96-well plate and fluorescence was immediately quantified at an excitation of 480 nm and an emission of 520 nm, using a Tecan Spark 10 M plate reader.

### Bacterial two hybrid (B2H) assays

Plasmid construction, cell cultivation, and experimental procedure of B2H assays were performed as described previously (Watzer et al. 2019, Selim et al. 2021 & 2023) on X-Gal plates supplemented with X-Gal (40 µg/mL), kanamycin (50 µg/mL), ampicillin (100 µg/mL), and IPTG (1 mM). We tested the N-terminal fusion of T25 subunit of Cya to SbtB was either N- or C-terminally fused to the T25 subunit of Cya, while the T18 subunit of Cya was fused either N- or C-terminally to the BCT1 subunit CmpC. The bicarbonate transporter BicA as well as the BCT1 subunit were only N-terminally fused to the T18 subunit of Cya. Primers used to generate fusion proteins are listed in (**Supplemental Table S4**). The leucine zipper interaction was used as positive control. As negative control the T25-SbtB fusion with an empty pUT18 vector was used. The *E. coli* BTH101 (Euromedex) was used for B2H assays. The B2H assays were performed at least three-times with three independent *E. coli* colonies to confirm the reproducibility and the specificity of the interactions.

## Data availability

The mass spectrometry proteomics data have been deposited to the ProteomeXchange Consortium via the PRIDE (Perez-Riverol et al. 2022) partner repository with the dataset identifier PXD045008.

## Acknowledgements

The project was funded by grants from the German Research Foundation (DFG) as part of the priority research program (SPP1879) to MHag (HA 2002/24-1) and to KF (Fo195/18-1), and by the Federal Ministry of Education and Research (BMBF) and the Baden-Württemberg Ministry of Science as part of the Excellence Strategy of the German Federal and State Governments to KAS (Projektförderung: PRO-SELIM-2022-14). KF and KAS acknowledge the infrastructural support by the Cluster of Excellence “Controlling Microbes to Fight Infections (CMFI)” (EXC 2124– 390838134). LC-MS/MS systems at the Department of Plant Physiology were supported through the DFG (grant No. INST 264/125-1 FUGG). Proteomics analysis was funded by DFG-FOR2816 (SCyCode). We are also grateful to Erik Zimmer (Tübingen University) for the excellent assistance in the pulldown experiments.

## Author contributions

KAS conceived, initiated, and supervised the research; KAS, MHaf, PS, and MHag designed research; MHaf, OM, and PS performed research; BM supervised the proteomics analysis; MHaf, OM, PS, and KAS analyzed data and prepared the figures; and MHaf and KAS wrote the manuscript with inputs from MHag, PS, BM and KF. All authors approved the final version of the manuscript.

## Competing interests

The authors declare no competing interest.

---

**Supplemental Fig. S1:**
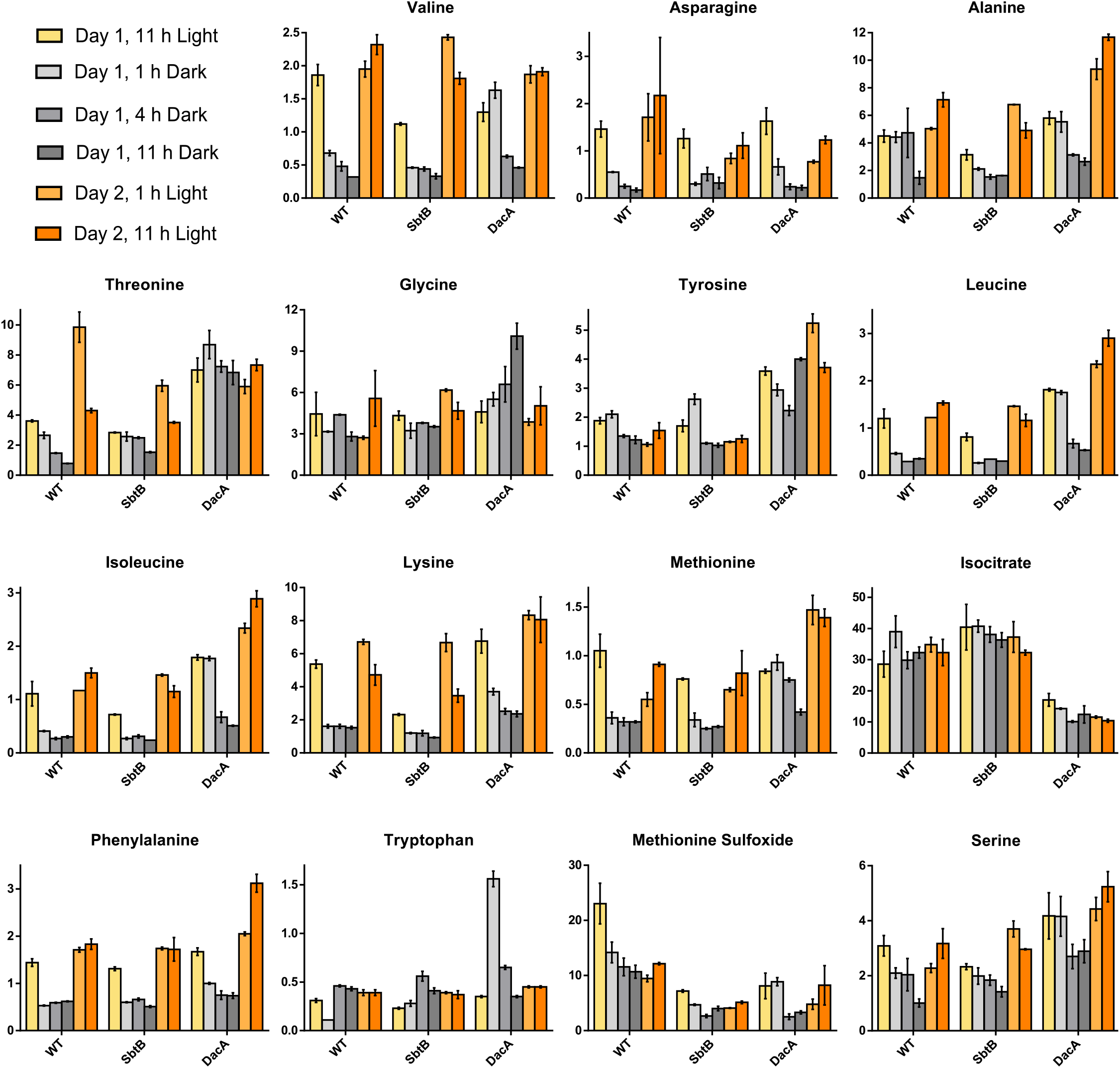
Targeted LC-MS based metabolomics analysis. Additional metabolites of interest are shown as ng ml^-1^ OD750^-1^, as indicated. Metabolites were quantified during 12 h day/12 h night diurnal rhythm, at 11 h of the first sampling day, 1, 4 and 11 h of the night-phase, and 1 and 11 h of the second sampling day, with color-code as indicated.

**Supplemental Fig. S2:**
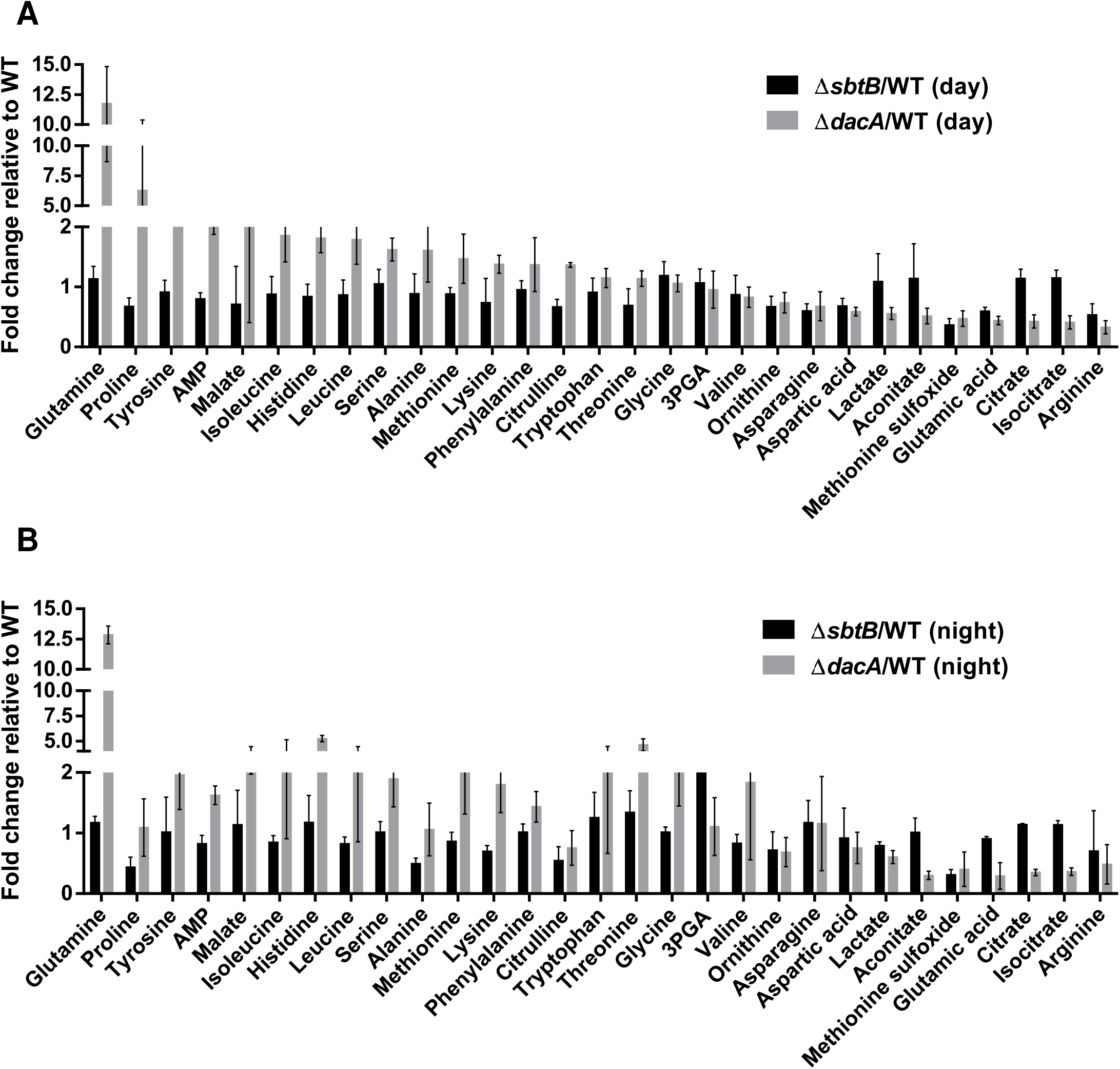
Relative fold-change of cellular metabolites within Δ*sbtB* and Δ*dacA Synechocystis* mutants compared to WT cells during diurnal rhythm. The Significant metabolic alterations are relative to wildtype (WT) metabolites (normalized to 1.0). **(A)** Average of identified cellular metabolites during the day phase. **(B)** Average of cellular metabolites during the night phase. For more details see (Supplemental Table S2).

**Supplemental Fig. S3:**
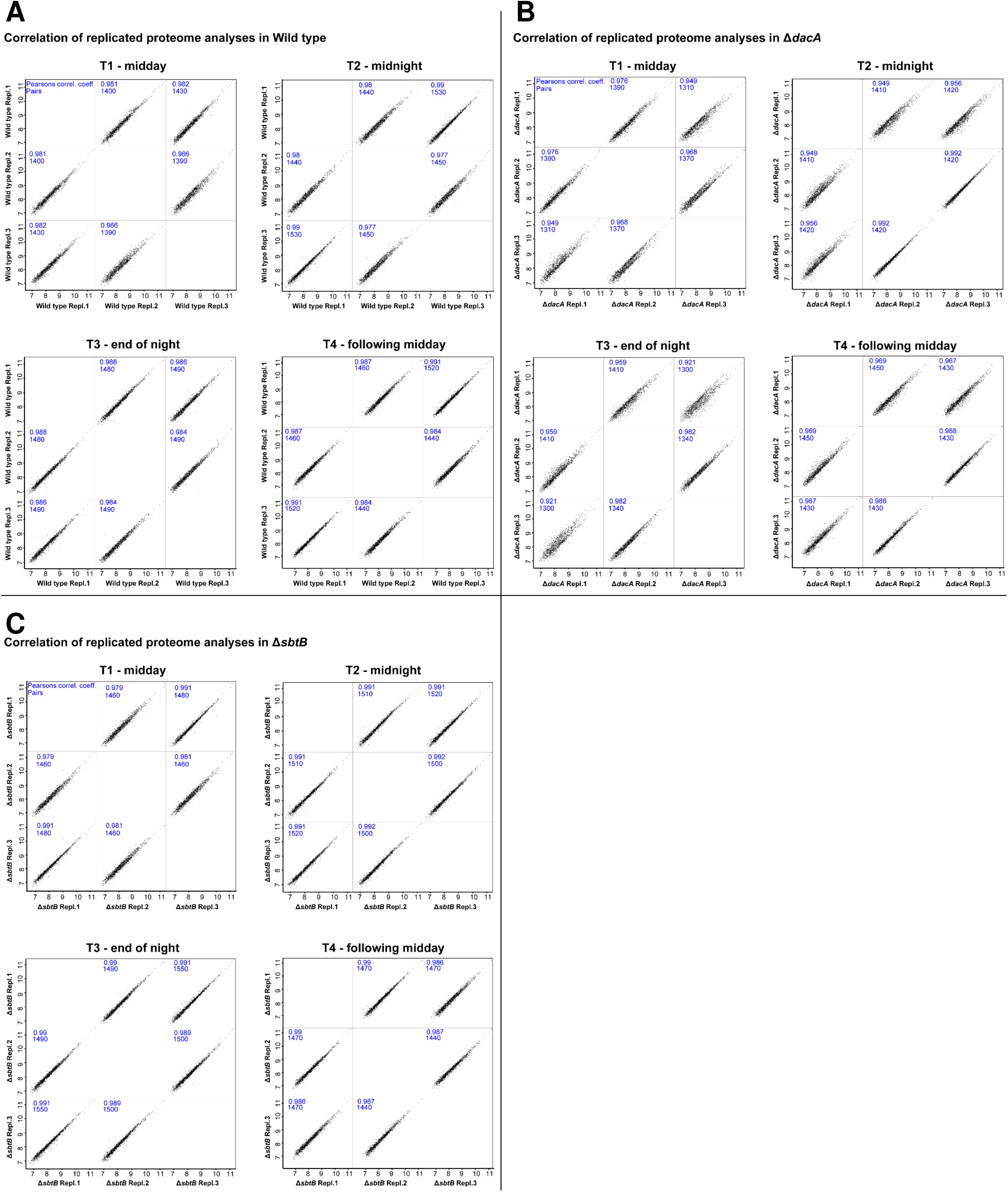
Intensity-based correlation of replicated proteome analyses. **(A-C)** Shown is the correlation of protein intensities (in Log_10_ scale) between independent replicates for each time point (T1-T4) and strain: **(A)** wildtype, **(B)** Δ*dacA* and **(C)** Δ*sbtB*. Pearson correlation coefficients are indicated for correlations between each two replicates as well as the number of matching proteins (rounded to tens).

**Supplemental Fig. S4:**
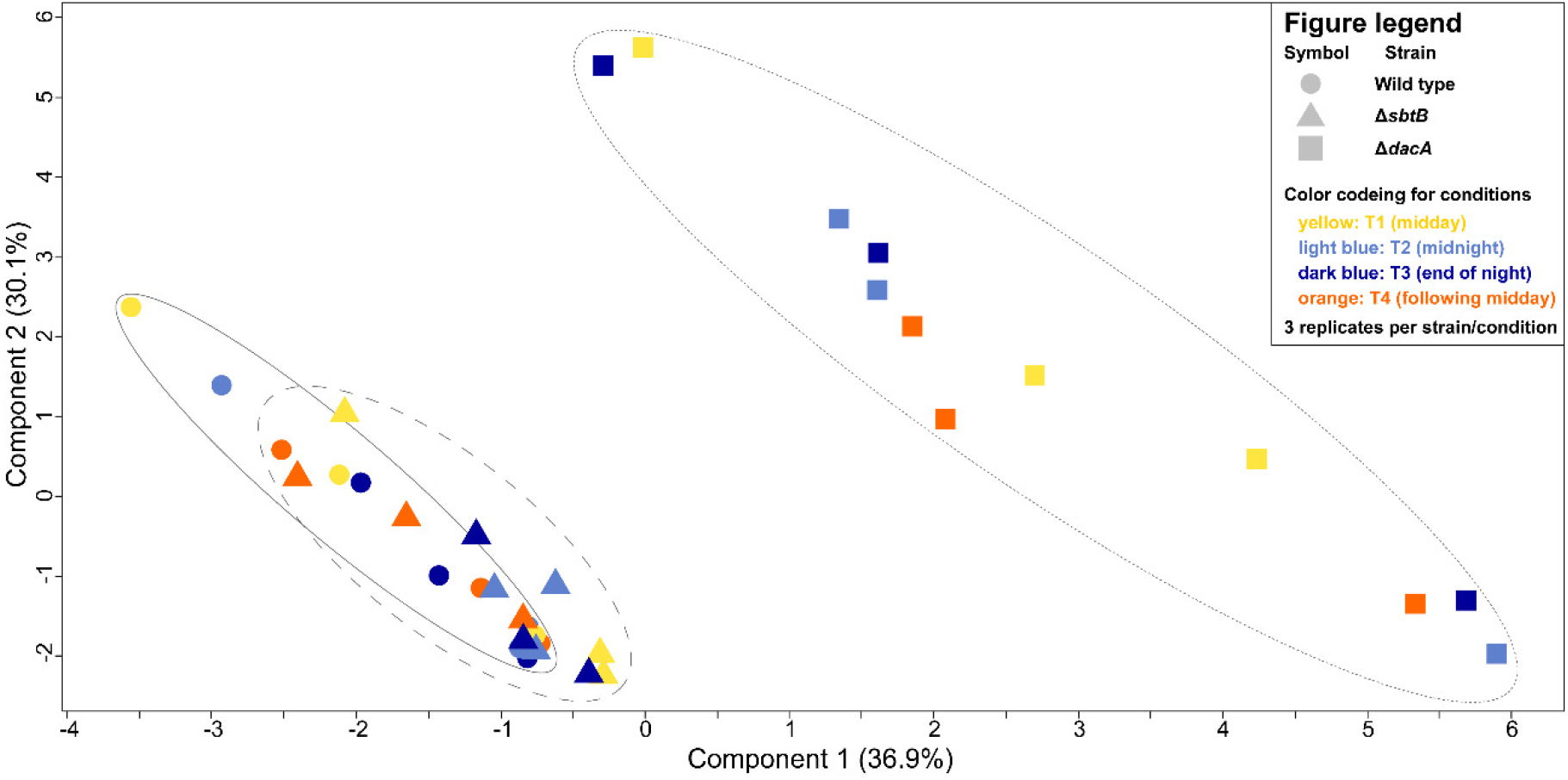
Proteomic landscape of Δ*dacA* in comparison to *Synechocystis* wildtype and Δ*sbtB* cells. Principal component analysis of protein abundance in the wild type (circles), Δ*sbtB* (triangles) and Δ*dacA* (squares) is shown at four different time points during the diurnal rhythm (color coding: T1 - yellow; T2 - light blue; T3 – dark blue; T4 - orange). Co-clustering of wildtype and Δ*sbtB* (overlay indicated by solid and dashed ellipses) indicates similar proteome compositions. Offset clustering of Δ*dacA* (dotted ellipse) indicates broad changes in protein abundance compared to wild type.

**Supplemental Fig. S5:**
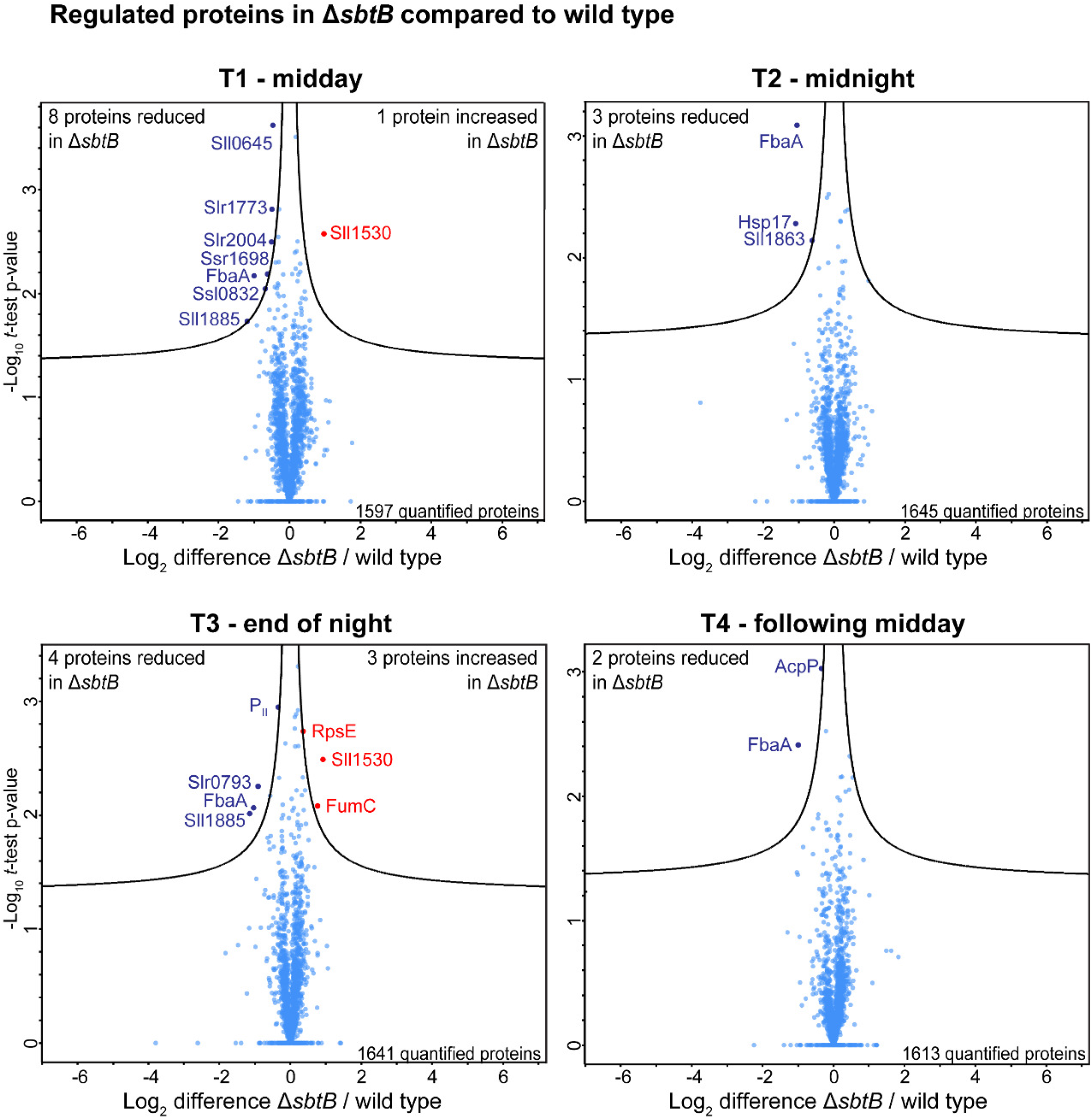
Proteome alterations of Δ*sbtB* mutnat. Quantitative comparison of protein abundance between Δ*sbtB* and wildtype at the four defined time points during the diurnal rhythm. Corresponding volcano plots indicate differences in protein abundance (Log_2_ difference Δ*sbtB*/wildtype) and corresponding −log_10_ p-values from *t*-test of three independent replicates per strain and time point. Proteins with significant changes in abundance at an p-value threshold of 0.05 are labeled in blue (reduced in Δ*sbtB*) and red (increased in Δ*sbtB*).

**Supplemental Fig. S6:**
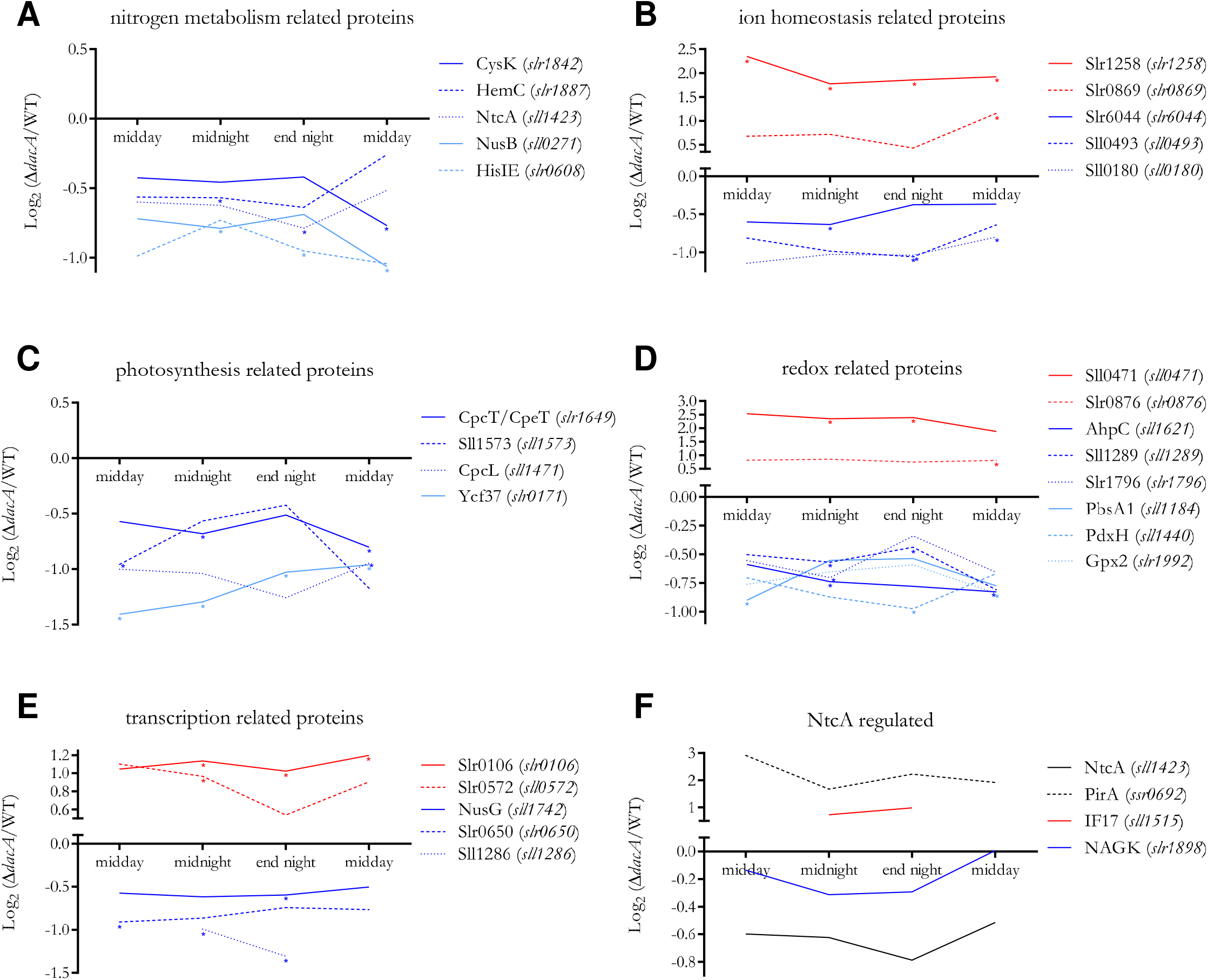
Proteomic landscape of Δ*dacA* in comparison to *Synechocystis* WT. **(A – E)** Log_2_ difference of Δ*dacA/*WT protein ratios (y-axis) throughout a diurnal cycle (x-axis) for exempler proteins involved in **(A)** nitrogen metabolism, **(B)** ion homeostasis, **(C)** photosynthesis and photosystem assembly, **(D)** redox regulation, and **(E)** transcriptional regulation; significant differences of Δ*dacA/*WT protein ratios are marked with asterisks. For more information (see Supplemental Table S3). **(F)** Proteomic landscape of exampler proteins under NtcA regulation in Log_2_ difference of Δ*dacA/*WT protein ratios (y-axis) throughout a diurnal cycle (x-axis).

**Supplemental Fig. S7:**
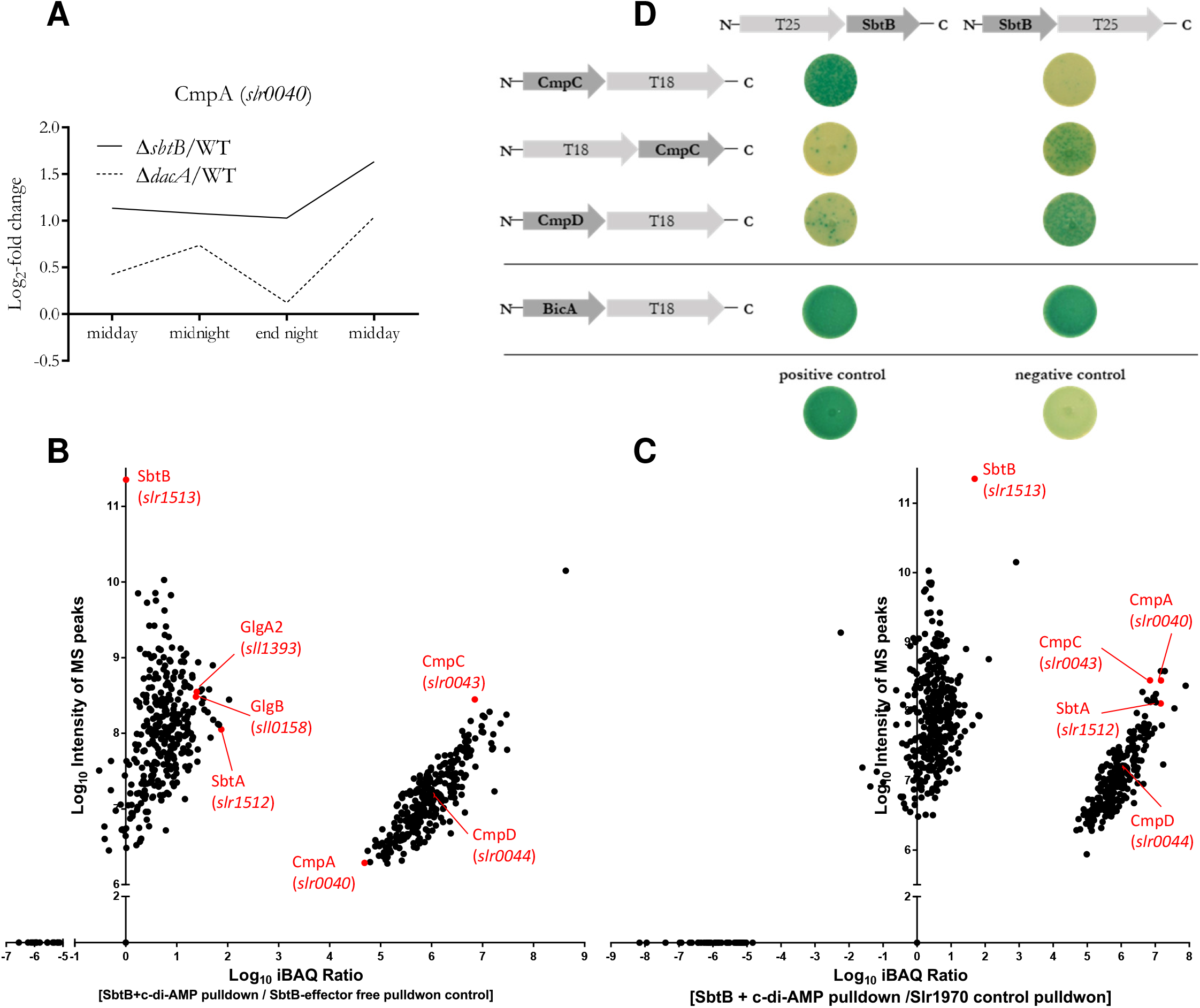
**(A)** Log2 difference of Δ*sbtB/*WT (solid line) and Δ*dacA/*WT (dashed line) protein ratios (y-axis) throughout a diurnal cycle (x-axis) for the subunit CmpA (Slr0040) of the bicarbnate uptak transporter complex CmpABCD, which is involved in substrate binding. **(B, C)** Identification of potential SbtB targets using Ni^2+^ magnetic beads-based pulldown with His_8_-tagged SbtB protein. The pulldowns were done using SbtB either in presence or absence of c-di-AMP (0.1 mM) (B) or using another unrelated His_8_-tagged Slr1970 protein as a control (C). Incubation of SbtB with c-di-AMP enriched the co-elution of different subunits of CmpABCD HCO_3_^-^ transporter complex, in practically CmpC with very high score of 289. While CmpA and CmpD subunits showed a moderate and low score of 152 and 22.5, respectively. The identification of known SbtB targets (SbtA and GlgB) validated our pulldown approach (Selim et al. 2018, 2021 & 2023). Eluates were analyzed by high accuracy LC-MS/MS to calculate protein enrichment ratios. The identified proteins were sorted by score and refined to remove unspecific binning proteins. Significantly enriched proteins were calculated based on Log10 of iBAQ values and plotted against the intensity of MS peaks of the identified/defined peptides. **(D)** Bacterial two hybrid (B2H) assay of the interaction of SbtB-T25/T25-SbtB with either CmpC-T18/T18-CmpC or CmpD-T18 on X-Gal supplemented LB agar (upper pannel). Bacterial two hybrid (B2H) assay of the interaction of SbtB-T25/T25-SbtB with BicA-T18 on either X-Gal supplemented LB agar (lower pannel). The N-terminal fusion of Cya-T25 subunit to SbtB together with N-terminal fusion of Cya-T18 subunit to either GlgB or GlgC, was used as positive and negative control, respectively (Selim et al. 2021 & 2023).

## Notes

### Competing Interest Statement

The authors have declared no competing interest.

